# Improving cell distribution on 3D additive manufactured scaffolds through engineered seeding media density and viscosity

**DOI:** 10.1101/815621

**Authors:** M. Cámara-Torres, R. Sinha, C. Mota, L. Moroni

## Abstract

In order to ensure the long-term *in vitro* and *in vivo* functionality of cell-seeded 3D scaffolds, an effective and reliable method to control cell seeding efficiency and distribution is crucial. Static seeding on 3D additive manufactured scaffolds made of synthetic polymers still remains challenging, as it often results in poor cell attachment, high cell sedimentation and non-uniform cell distribution, due to gravity and to the intrinsic macroporosity and surface chemical properties of the scaffolds. In this study, the bio-inert macromolecules dextran and Ficoll were used for the first time as temporary supplements to alter the viscosity and density of the seeding media, respectively, and improve the static seeding output. The addition of these macromolecules drastically reduced the cell sedimentation velocities, allowing for homogeneous cell attachment to the scaffold filaments. Both dextran- and Ficoll-based seeding methods supported human mesenchymal stromal cells viability and osteogenic differentiation post-seeding. Interestingly, the improved cell distribution led to increased matrix production and mineralization compared to scaffolds seeded by conventional static method. These results suggest a simple and universal method for an efficient seeding of 3D additive manufactured scaffolds, independent of their material and geometrical properties, and applicable for bone and various other tissue regeneration.

## 1. Introduction

*In vivo*, cells are surrounded by a complex 3D microenvironment that supports their growth and assembly into functional tissues. Upon tissue damage caused by trauma or disease, the structural integrity of this extracellular matrix (ECM) can be compromised leading to lack of essential cues for functional tissue performance, and the requirement of suitable replacements. By using scaffolds, tissue engineering and regenerative medicine (TERM) aim to provide artificial ECM-like structures, which try to mimic the native tissue in terms of architecture, chemistry or mechanical properties to guide cells into the first steps of the regeneration process [1]. To allow cell infiltration, tissue ingrowth and vascularization, 3D scaffolds with a highly interconnected porous structure are required [2, 3]. In the past decade, additive manufacturing and, more specifically, melt extrusion-based techniques, have emerged as one of the most appealing technologies to produce scaffolds for TERM [4, 5]. Compared to other conventional methods such as gas foaming/particulate leaching [6, 7], freeze-drying [8, 9] or phase separation [10, 11], additive manufacturing allows the reproducible fabrication of complex 3D structures with interconnected, high porosity, from various types of biocompatible and biodegradable polymers with optimum mechanical properties desired for tissue regeneration applications.

Efficient cell seeding of 3D scaffolds is a key step towards the development of *in vitro* tissues for TERM applications. Cell density and their spatial distribution in macroporous scaffolds are critical to ensure the functionality of the engineered tissue, as these parameters will dictate intercellular and cell-material interactions, thereby affecting different cellular behaviors such as migration, proliferation, differentiation and matrix production [12, 13]. To improve cell distribution within 3D scaffolds, different dynamic seeding methods have been applied. Compared to static seeding, dynamic seeding methods, which apply an external force to facilitate cell penetration, such as flow perfusion, centrifugation, orbital shaking, and spinner flask, have shown to result in better cell distribution [14-18]. However, due to the intrinsic scaffold characteristics, i.e. large and interconnected pore architectures, most of the dynamic seeding techniques show limited applicability for seeding additive manufactured (AM) scaffolds. Furthermore, the lack of specific cell adhesion sites in the majority of the polymers suitable for AM leads to poor cell–material interactions, which along with the gravitational force results in cell sedimentation during static seeding and, therefore, non-uniform cell distribution within the scaffold. Nevertheless, the simplicity of the static seeding over dynamic seeding has led to further developments of this method for seeding AM 3D scaffolds, starting by the modification of basic seeding parameters such as cell number, seeding volume, seeding time and seeding vessel optimization. For instance, seeding times up to 4h, seeding volumes comparable to the scaffold’s free volume and not very large cell numbers have shown to give the highest cell seeding efficiency upon static seeding [19-21]. On the other hand, different surface modification techniques capable of improving the chemical and biological performance of synthetic polymers have also shown to increase cell adhesion in AM 3D scaffolds. For example, plasma modification using oxygen [22] or nitrogen [23, 24] have shown to increase cell adhesion of osteoblastic cell lines in poly(*ε*-caprolactone) (PCL) scaffolds. Chondrocytes adhesion on poly(ethylene oxide terephthalate)/poly(butylene terephthalate) (PEOT/PBT) AM 3D scaffolds has also been improved by the addition of carboxylic groups to the polymer surface via acrylic acid plasma polymerization [25]. Surface functionalization using adhesive polymers such as poly(dopamine) have also demonstrated to improve adhesion of different cell types on PCL and poly(lactic acid) (PLA) scaffolds [26-28]. As an alternative to chemical modification of polymer surfaces, better cell attachment along the scaffold cross section has been attained via decreasing the flow rate of the seeding media through the scaffold’s pores [29, 30]. By varying the strand distance or the angle of deposition in each layer, AM 3D scaffolds with pore size gradients along the vertical direction [29] or with different lay-down patterns [30] were fabricated and resulted in enhanced cell attachment and distribution. This was, however, in exchange of varying the scaffolds mechanical properties and, more importantly, reducing the pore size, which hinders cell migration, nutrients and oxygen availability, waste management, and angiogenesis for pore sizes smaller than 300 *µ*m [3, 31-33].

A large improvement in cell seeding efficiency and distribution was recently obtained by seeding AM 3D scaffolds with cell-laden hydrogels, which showed to retain almost the complete cell number mixed within the hydrogel precursor [34-38]. However, these hybrid systems are multi-step and require fast gelation times. In addition, cells are exposed to two different microenvironments: the hard scaffold filaments and the soft hydrogel carrier. As it is widely accepted that substrate stiffness can direct cell fate [39], having these types of dual systems might mask the effect of the scaffold material and lead to unexpected results.

Although previous strategies have shown some improvement on cell seeding efficiency and distribution, they require modification of the scaffold’s surface chemistry or architecture, influencing the overall microenvironment. Therefore, there is still lack of methodology and knowledge of an elegant cell seeding method applicable to all AM 3D scaffolds regardless of their characteristics. Here, we introduce a novel seeding technique based on controlling cell settling velocity upon static seeding that can be applied to all types of AM 3D scaffolds without requiring any chemical or architectural modification. Two bioinert macromolecules (macroMs) were employed for the first time as supplements to independently tune i) the viscosity or ii) the density of the cell seeding media. Cells seeded with this novel technique were morphologically characterized using immunofluorescence and scanning electron microscopy. Furthermore, the osteogenic differentiation of human mesenchymal stromal cells (hMSCs) seeded with the macroM-based solutions on AM 3D scaffolds was investigated using various biochemical and molecular assays to evaluate the effect of macroMs on a specific cell differentiation.

## 2. Materials and methods

### 2.1. Scaffolds fabrication

Scaffolds were fabricated via a melt-extrusion based AM technique (Bioscaffolder 3.1, SysENG, Germany). The copolymer poly(ethylene oxide terephthalate)/poly(butylene terephthalate) (PEOT/PBT), with a PEO molecular weight of 300 kDa and PEOT/PBT blocks ratio 55/45 wt% (300PEOT55PBT45, PolyVation, The Netherlands) was used. Briefly, the cartridge was filled with PEOT/PBT pellets, heated at 195 °C and extruded by applying a pressure of 4 bar, an auger screw rotation of 40 rpm and a translation speed of 10 mm*s^-1^. The scaffold architecture consisted of a 0-90 pattern, with a 250 *µ*m fiber diameter, 200 *µ*m layer thickness and 750 *µ*m strand distance (center to center), giving an expected x-y porosity of approximately 500 *µ*m. Cylindrical scaffolds of 4 mm diameter and 4 mm height were punched out from a 20×20×4 mm manufactured block using a biopsy punch and used for further experiments.

### 2.2. Density and viscosity measurements of Dextran and Ficoll based solutions

Ficoll-Paque™ Plus solution (GE Healthcare) was diluted to 80, 60, 40 and 20 vol% with cell culture medium (CM), consisting of αMEM with Glutamax and no nucleosides (Gibco) supplemented with 10 vol% FBS (Sigma-Aldrich), penicillin (100 U/ml) and streptomycin (100 *µ*g/ml) (Fisher-Scientific). Similarly, 10, 5, and 2.5 wt% dextran (500 kDa, Pharmacosmos) solutions were prepared in CM. Densities were calculated by weighing known volumes of each of the solutions (accuracy ± 0.1 ml) in a high precision balance (accuracy ± 0.1 mg) and applying the density formula.. The relative viscosity (with respect to water) of the solutions was empirically determined by timing the flow of water and the same volume of each solution through the same fluidic circuit at the same constant pressure.

### 2.3. Determination of hMSCs density using density gradient separation

Density of hMSCs was measured using Ficoll-Paque™ Plus (Ficoll) density gradient centrifugation. The Ficoll solution was diluted to 80, 60, 40 and 20 vol% with appropriate volumes of CM containing phenol red, which helped to visualize the layers of the gradient. Starting from 100 vol% Ficoll solution, 2ml of each solution were carefully layered on top of each other in decreasing order in a 15 ml conical tube. One million cells in CM were layered on top of the 20% Ficoll layer. The tube was then centrifuged at 500 rcf for 20 min at room temperature (RT). Subsequently, the layers were carefully separated by pipetting and the cells in each layer were imaged via brightfield microscopy and quantified using a Neubauer chamber. As a comparative method, cell density was additionally determined through buoyancy force-driven separation, which does not require the application of mechanical forces. For the buoyancy separation, the various concentrations of Ficoll media were similarly layered in a 15 ml conical tube starting from 100 vol% Ficoll solution. Here, 1 million cells in medium were carefully pipetted at the bottom of the tube under the 100% Ficoll layer using a long syringe needle, without disturbing the layers. The tube was then incubated at RT for 3h to let cells be pushed by buoyancy forces to layers of similar density. After separation, the layers were collected separately and cells present in each solution were imaged and quantified using a Neubauer chamber.

### 2.4. Cell seeding and culture

#### 2.4.1. Cell expansion

HMSCs isolated from bone marrow were purchased from Texas A&M Health Science Center, College of Medicine, Institute for Regenerative Medicine (Donor d8011L, female, age 22). Cryopreserved vials at passage 3 were plated at a density of 1,000 cells*cm^-2^ in tissue culture flasks and expanded until approximately 80% confluency in CM without Pen/Strep at 37 °C / 5% CO_2_.

#### 2.4.2. Cell seeding and culture in 3D scaffolds

Scaffolds were disinfected in 70% ethanol for 20 min, washed 3 times with phosphate buffered saline (PBS) and incubated overnight in CM for protein attachment. Before seeding, scaffolds were dried on top of a sterile filter paper and placed in the wells of a non-treated well plate. Trypsinized passage 4 hMSCs suspension was centrifuged for 5 min at 500 rcf and the cells were then resuspended in CM, Dextran-based CM, or Ficoll-based CM at a density of 200,000 cells per 37 *µ*l. Dextran-based CM consisted of 10 wt% dextran in CM and Ficoll-based CM consisted of 60 vol% Ficoll in a FBS adjusted cell culture medium (α-MEM supplemented with 25 vol% FBS and 1 vol% Penstrep, to obtain 10 vol% FBS in the final medium). A 37 *µ*l droplet of each cell suspension each was placed on top of each scaffold and incubated for 4 h at 37 °C / 5% CO_2_ to allow cell attachment. After this time, scaffolds were transferred to new wells containing 1.5 ml of basic media (BM) (CM supplemented with 200 μM L-Ascorbic acid 2-phosphate (Sigma-Aldrich)). BM was replaced after 24h and every two or three days from then on. To evaluate hMSCs osteogenic differentiation, scaffolds were cultured for another 21 days in BM or mineralization media (MM) after 7 days in BM. MM consisted of BM supplemented with 10 nM dexamethasone (Sigma-Aldrich) and 10 mM β-glycerophosphate (Sigma-Aldrich).

To visualize the potential entrapment of macromolecules within the scaffold after seeding, cells were resuspended in 10 wt% dextran-FITC (Fluorescein Isothiocyanate-Dextran 500 kDA, Sigma-Aldrich) or 60 vol% Ficoll-FITC at the same cell density (200,000 cells in 37 *µ*l). The Ficoll-FITC based CM was prepared by first dissolving 5.7% (wt/vol) polysucrose-FITC (400 kDa, Sigma-Aldrich) and 9% (wt/vol) sodium diatrizoate hydrate (Sigma-Aldrich) in sterile water, to match Ficoll-Paque™ Plus composition. 60 vol% of this solution was prepared in FBS adjusted cell CM as previously described. A 37 µl droplet of each cell suspension was placed on top of each scaffold and incubated for 4 h at 37 °C/ 5% CO_2_ to allow cells attachment. After this time, scaffolds were transferred to new wells containing 1.5 ml of BM and cultured for 24h.

To rule out macromolecular adhesion as the cause of improved cell seeding, scaffolds were pre-incubated for 4h with dextran- and Ficoll-based CM. Non-adsorbed macromolecules were removed by a single scaffold wash with PBS and the scaffolds were then dried on top of a sterile filter paper. Subsequently, scaffolds were seeded with 37 µl of CM containing 200,000 cells and incubated for 4 h at 37 °C / 5% CO_2_ to allow cell attachment, after which scaffolds were analyzed.

#### 2.4.3. Cell seeding in 2D for viability and osteogenic differentiation evaluation

Trypsinized passage 4 cells were centrifuged and resuspended in CM, dextran-based CM or Ficoll-based CM at a density of 200,000 cells per 37 µl. Cell suspensions were incubated at 37 °C/ 5% CO_2_ in plates mimicking the 4h seeding in 3D scaffolds in contact with the macroMs. To assess the viability of cells when in a macroMs solution, incubation was done in presence of NucBlue® Live and NucGreen® Dead reagents (ReadyProbes® Cell Viability Imaging Kit, Invitrogen) as defined by the manufacturer protocol and imaged at t = 0 h and t = 4 h after seeding using an inverted fluorescent microscope (Eclipse, Ti2-e, NIKON). For optimal visualization in the reported images, live cells were false colored in green while dead cells were given a red color. After 4 h, the remaining cell suspensions (without cell viability reagents) were centrifuged, the macroMs containing media was removed and cells were resuspended in fresh CM to mimic the post-seeding culture condition of cells in scaffolds. Cells were counted and seeded at a density of 5,000 cells*cm^-2^ in tissue culture polystyrene well plates. Media was refreshed every two or three days. To evaluate hMSCs osteogenic differentiation, scaffolds were cultured for 21 days in BM or MM after 7 days in BM.

### 2.5 Imaging and quantification of cell distribution within scaffold cross section

After seeding and 4h incubation for attachment, scaffolds were washed with PBS and fixed with 4 wt% paraformaldehyde for 30□min, followed by washing steps with PBS. Cells were permeabilized using 0.1 vol% Triton-X for 30□min, washed twice with PBS and incubated with phalloidin (Alexa Fluor 568, 1:75 dilution in PBS) for 1h at RT. Finally, samples were washed with PBS. The bottom and cross section of scaffolds were imaged using an inverted fluorescent microscope. Cross section images were analyzed using ImageJ software. Briefly, images were converted to 8-bits format and the local contrast was enhanced (blocksize 50, maximum slope 3, no mask). Next, the background was subtracted, using a rolling ball radius of 400 pixels, and the images were converted to binary (rendering regions with cells in white and the rest in black). Each image was divided into four regions corresponding to heights of 0-1 mm, 1-2 mm, 2-3 mm, 3-4 mm starting from the bottom of the scaffolds and the total amount of white pixels in each region was quantified and normalized to the total number of pixels in the scaffold cross section. Values are reported as normalized cell coverage area in percentage.

### 2.6. Imaging of cell viability and macromolecules entrapment in 3D scaffold

After 4h seeding and 1 and 7 days of culture, dead cells were stained for 20 minutes prior to fixation using the LIVE/DEAD™ Fixable Dead Cell Stain Kit (Thermo Fisher Scientific) at a concentration of 0.5 µl stain in 500 µl Hank’s Balanced Salt Solution per scaffold. Subsequently, samples were washed and fixed with 4 wt% paraformaldehyde for 30□min, followed by three washing steps with PBS. Cells were stained with phalloidin as stated in section 2.5. Live/dead images of scaffolds at day 1 and 7 were acquired using an inverted fluorescent microscopy. Confocal laser scanning microscopy of macroMs entrapment/dead cells at 4h and day 1 was performed with a Tandem confocal system (Leica TCS SP8 STED), equipped with a white light laser (WLL). Samples were excited with the dye specific wavelengths using the WLL or a photodiode 405 in the case of DAPI. Emission was detected with PMT detectors (DAPI) or HyD detectors (phalloidin, dextran-FITC, Ficoll-FITC, dead cells).

### 2.7. Biochemical assays

#### 2.7.1. ALP assay

ALP activity was evaluated after 7 and 21 days of culture in MM (time points day 14 and 28, respectively). 3D scaffolds were collected at every time point and washed with PBS, cut in half, stored at −80 °C and freeze-thawed 3 times. Samples were incubated for 1h at RT in a cell lysis buffer composed of 0.1 M KH_2_PO_4_, 0.1 M K_2_HPO_4_, and 0.1 vol% Triton X-100, at pH 7.8. 40 µl of the chemiluminescent substrate for alkaline phosphatase CPD star-ready to use reagent was added to 10 µl of cell lysate. After 15 min incubation, luminescence (emission= 470 nm) was measured using a spectrophotometer (Biodrop). Remaining cell lysates were used for DNA quantification. Values are reported normalized to DNA content.

#### 2.7.2. DNA assay

DNA assay was performed on cells cultured on 3D scaffolds after 4h of seeding (time point 4h), after 7 days in BM (time point day 7), and after an extra 7 and 21 days of culture in BM or MM (time point day 14 and 28, respectively). In addition, DNA quantification was also performed on 2D samples using CyQUANT Cell Proliferation Assay Kit (Thermo Fisher Scientific). Samples (lysed samples from ALP activity assay or frozen samples after time point collection) were incubated overnight at 56 °C in Proteinase K solution (1mg*ml^-1^ Proteinase K (Sigma-Aldrich) in Tris/EDTA) for matrix degradation and cell lysis. Subsequently, samples were freeze-thawed three times and incubated 1h at RT with lysis buffer (cell lysis buffer from the kit diluted 20x in dH_2_O) containing RNase A (1:500) to degrade cellular RNA. Lysed samples were incubated with the fluorescent dye provided by the Cyquant kit (1:1) for 15 min and fluorescence was measured (emission/excitation = 520/480 nm) with a spectrophotometer. DNA concentrations were calculated from a DNA standard curve.

#### 2.7.3. Alizarin red S staining and quantification

Calcium mineralization was qualitatively determined by alizarin red S staining after 7 and 21 days of culture in BM or MM (time point day 14 and 28, respectively). 3D scaffolds were washed with PBS and fixed with 4 wt% paraformaldehyde for 30□min, followed by three washing steps in distilled water. Subsequently, scaffolds were cut in half and each section was stained with alizarin red S solution (60 mM, pH 4.1-4.3) for 20 min at RT, and washed several times with distilled water until no more stain was leaching out. Images were taken using a stereomicroscope (Nikon SMZ25).

After imaging, a protocol adapted from Gregory et al. [40] was used to quantify calcium deposition. Briefly, stained samples were incubated in Eppendorf’s for 1h at RT with 30 vol% acetic acid while shaking, followed by 10 min incubation at 85 ^°^C. Afterwards, scaffolds were removed and solutions were centrifuged at 20,000 rcf for 10 min. An appropriate volume of 5M ammonium hydroxide was added to the supernatants to readjust the pH to 4.1-4.3. The absorbance was measured at 405 nm using a spectrophotometer. Concentration of alizarin red was calculated from an alizarin red standard curve and the values were normalized to DNA content.

### 2.8. Gene expression

Gene expression analysis was performed at day 14 (7 days in MM) and day 28 (21 days in MM). RNA was extracted from cells in scaffolds using a Trizol isolation method at the selected time points. Briefly, samples were transferred to Eppendorf tubes and Trizol was added. ECM was precipitated at the bottom of the tube through a centrifugation step at 12,000 rcf for 5 min. The supernatant was transferred to a new tube and chloroform was added to isolate the RNA, present in the aqueous phase after phase separation via centrifugation at 12,000 rcf for 5 min. RNA was further purified using RNeasy mini kit column (Qiagen), according to the manufacturer’s protocol. The purity and quantity of total RNA was evaluated using a spectrophotometer. Reverse transcription was performed using iScript™ (Bio-Rad) following suppliers’ protocol. The obtained cDNA was used in combination with SYBRGreen master mix (Qiagen) and the selected primers (Table S1, Supplementary Information) to perform qPCR using CFX Connect™ Real-Time System (Bio-Rad) under the following conditions: cDNA was denaturated for 3 min at 95 °C, followed by 40 cycles consisting of 15 s at 95 °C and 30 s at 65 °C. Additionally, a melting curve was generated for each reaction in order to test primer dimer formation and non-specific amplification. Gene transcription was normalized to the transcription ofthe housekeeping human B2M gene. The 2^-ΔΔCt^ method was used to calculate relative gene expression for each target gene. Normalization was done with respect to relative gene expression of cells in conventional seeding (CS) scaffolds at day 14.

### 2.9. Statistical analysis

Analysis of statistics was conducted with GraphPad Prism (version 8.0.1). A one-way or two-way ANOVA was performed with a post-hoc comparison to evaluate statistical significance. Data is shown as average of minimum 3 independent replicas with error bars indicating the standard deviation.

## 3. Results and discussion

### 3.1. Optimizing macromolecular seeding methods

When cells are seeded on a 2D substrate, gravity acts as one of the main parameters favoring cell attachment. On the contrary, gravity acts as a counterforce to cell attachment during static seeding of 3D scaffolds fabricated from bioinert polymers and with large and interconnected pores, where cell attachment is already dramatically hindered because of lack of adhesion motifs. Rather than modifying the scaffold material properties or architecture, in this study we tackled the cell sedimentation problem from a different perspective, the cell seeding media properties. This is particularly a beneficial approach as it can be applied to all AM 3D scaffolds independent on their chemistry or geometries. By controlling the physical properties of the seeding media (viscosity and density) cell settling velocity can be appropriately reduced, allowing cells to float longer in the suspension, thereby giving them more time to attach, and hence promoting homogeneous cell attachment on the scaffold filaments (Error! Reference source not found.). If cells in suspension are considered as rigid spheres, upon which frictional (viscosity related), buoyant (density related) and gravitational forces are exerted, their settling velocity in medium can be derived from the Stokes law [41]. When the sum of the frictional and buoyancy forces exerted on the cell, due to the surrounding medium, balance the gravitational force, the constant settling velocity (v) can be defined by:

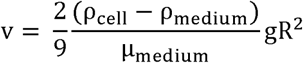

Where *ρ*_cell_ refers to cell density, *ρ*_medium_ *ρ*medium refers to density of the surrounding cell culture medium, µ_medium_ refers to the viscosity of the surrounding cell culture medium, g is the gravitational force, and R the cell hydrodynamic radius. According to this equation, to maintain cells in suspension the settling velocity can be decreased by increasing the media viscosity or canceled by matching the density of the media and the cells. In this study, we investigated both approaches independently. The media viscosity has been increased with the addition of dextran. Dextran is an inert natural biocompatible macromolecule that has been widely used in the biomedical field. It has shown to promote ECM deposition *in vitro* not only when used as macromolecular crowder mimicking the highly crowded/dense native ECM [42-44], but also when employed as media thickener to increase fluid shear forces in perfusion culture [45]. It has also been used to study erythrocyte aggregation and blood viscosity changes [46-48]. As shown in **Fig. S1**A (Supplementary Information), by the addition of dextran up to 10 wt%, the viscosity of CM increases exponentially up around 25 times. As our second approach, the density of the seeding medium has been increased by the addition of Ficoll-Paque™ Plus to match the cell density and therefore obtain a theoretical settling velocity equal to zero (v=0). Ficoll-Paque™ Plus (Ficoll) is a Ficoll aqueous solution that contains sodium diatrizoate, a compound that allows to have a high density solution while maintaining its viscosity relatively low (**Fig. S1**B, Supplementary Information). Notably, the viscosity of this solution is only ∼3 times higher than CM viscosity. Ficoll is a highly branched polysaccharide that has been used for the isolation of cells from blood samples by centrifugation [49, 50], as well as macromolecular crowding agent [44, 51, 52]. A density gradient of Ficoll (**Fig. S1**D, Supplementary Information) allowed for the experimental determination of the density of the hMSCs, which is known to be lower than 1.073 g*ml^-1^ [53, 54]. Both separation by gradient centrifugation and by buoyancy forces determined that the density of the population of hMSCs lied in between 1.034 and 1.052 g*ml^-1^ corresponding to 40 vol% and 60 vol% Ficoll in CM (**Fig. S2**, Supplementary Information). In order to encompass the majority of the hMSCs population (i.e. maintain most of the cells in suspension) 60 vol% Ficoll solution (Ficoll-based CM) was chosen as the density-based media for seeding 3D scaffolds, with a relatively ignorable influence on the viscosity of CM (only 1.65 fold increase) (**Fig. S1**B, Supplementary Information). On the other hand, the viscosity is considered the only effective parameter on the cell settling velocity in a 10 wt% Dextran solution (Dextran-based CM), which was chosen as the viscosity-based media for seeding the scaffolds. 10 wt% dextran density (1.024 g*ml^-1^) is significantly lower than 60 vol% Ficoll density (1.052 g*ml^-1^), and therefore, than hMSCs density. Specifically, its density is comparable to that of a 20 vol% Ficoll solution (1.027 g*ml^-1^) (**Fig. S1**C and E, Supplementary Information), upon which cells were observed to sediment (**Fig. S2**, Supplementary Information). Therefore, given that viscosity and density are the effective parameters on cell settling velocity of the chosen dextran and Ficoll based solutions respectively, these two methods are considered independent approaches to tackle the cell sedimentation problem upon scaffolds static seeding, from here on referred as MS-Dextran and MS-Ficoll, respectively.

### 3.2. Cell distribution using MS methods

As it was expected, cells seeded on highly porous PEOT/PBT scaffolds with the CS method sedimented at a theoretical velocity of 1.8 µm*s^-1^ (**Fig. S1**F, Supplementary Information) and after 4h post seeding most were found in the lower region of the scaffolds, as well as attached to the bottom (**Fig. 2**A). This is due to the lack of cell adhesion motifs on the PEOT/PBT filaments not allowing for rapid cell attachment while the cells settle under gravity, due to the fast flow velocity and to the scaffold’s pore size. Consistent with a previously published report, a cell monolayer was formed at the bottom of scaffolds seeded with CS method [21], which is likely due to the high cell density enabling cell-to-cell and cell-to-scaffold contact after their fast sedimentation.

**Fig. 1.**
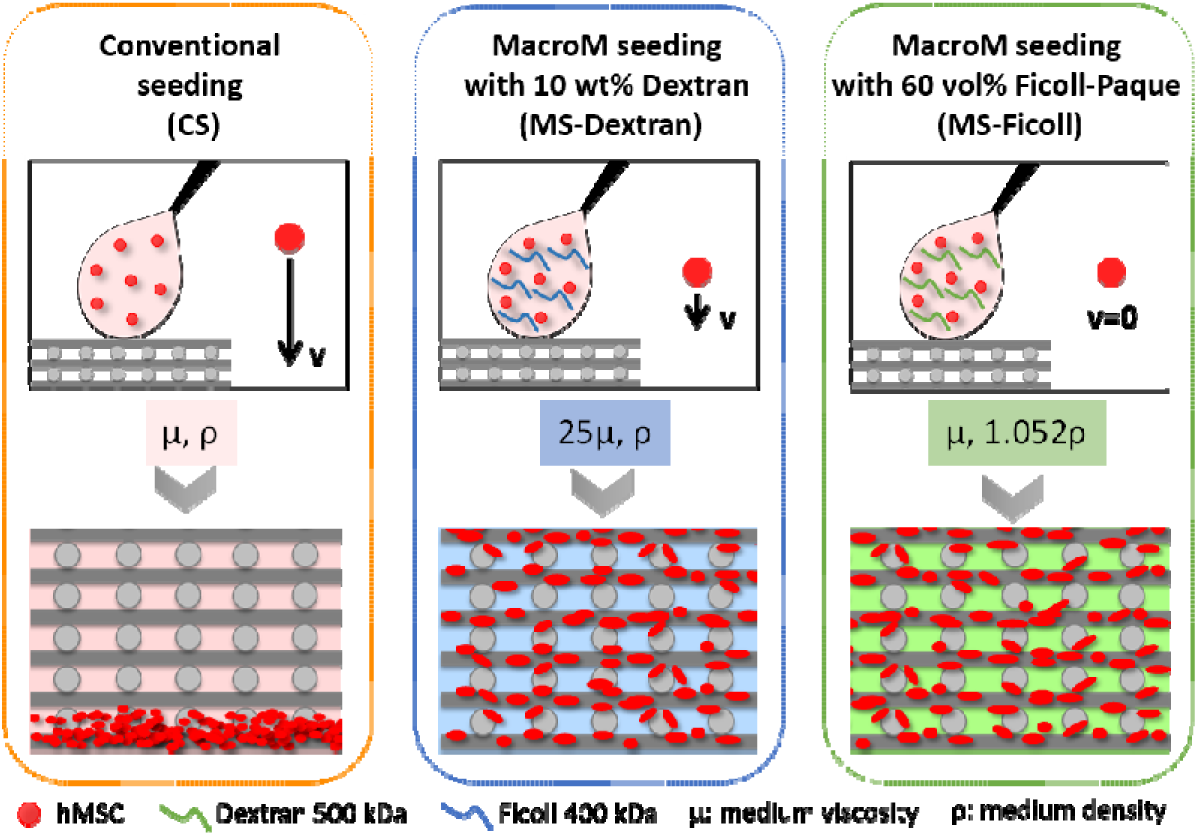
Schematic representation of the static seeding approaches discussed in this study: Conventional seeding (CS), Macromolecular seeding with 10 wt% dextran (MS-Dextran) and macromolecular seding with 60 vol% Ficoll-Paque (MS-Ficoll). Influence of the properties of the cell seeding media (viscosity and density) on the settling velocity of cells and, therefore, on cell attachment along scaffold height is depicted.

**Fig. 2.**
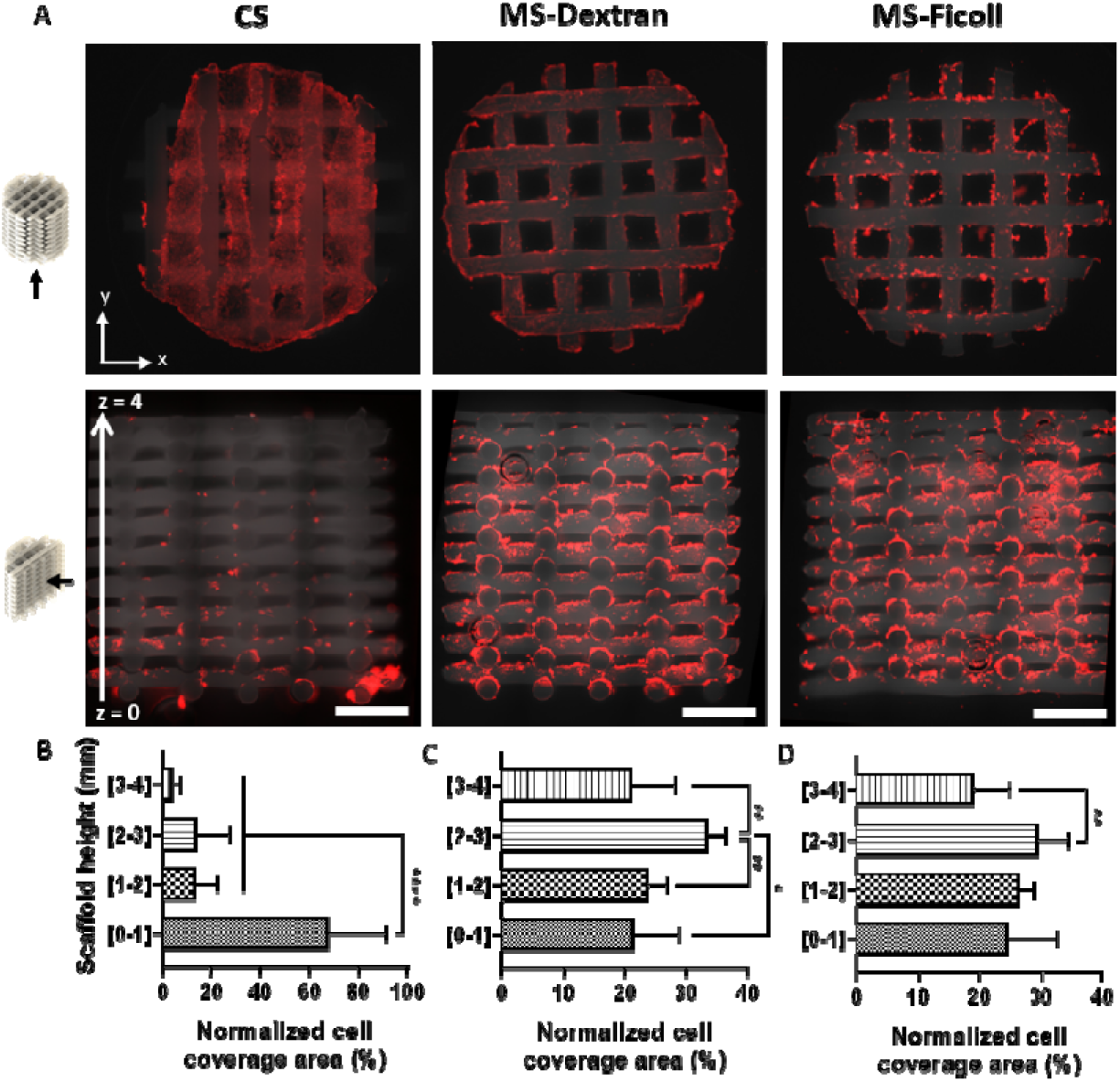
Fluorescent staining (F-actin, red) showing that MS methods improved cell distribution along scaffold cross section. (a) Fluorescence microscopy images of hMSCs in the bottom sides (top) and cross sections (bottom) of scaffolds seeded with the CS, MS-Dextran and MS-Ficoll methods and (b-d) respective quantification of cell distribution across the scaffold height. Statistical significance performed using one-way ANOVA with Tukey’s multiple comparison test (* p < 0.05; ** p < 0.01; **** p < 0.0001). Scale bars 1mm.

To assess the cell distribution across the scaffolds quantitatively, their cross section images were divided into four horizontal regions of 1 mm each, from 0 to 4 mm height. As demonstrated in **Fig. 2B**, around 70% of the total cell number found in the scaffold cross section is concentrated in the lowest 1 mm region of the scaffold when seeded with the CS method. This is consistent with the calculations derived from the settling velocity, according to which cells displaced 2.6 mm (∼75% of the scaffold height) towards the bottom of the scaffold during the 4h seeding. Importantly, theoretical cell settling velocity decreased ∼50 fold for MS-Dextran when compared with the CS method, or remained zero when seeded with the MS-Ficoll method, resulting in a theoretical displacement of ∼55 µm or 0 µm towards the bottom of the scaffold during the 4 h seeding, respectively (**Fig. S1**F, Supplementary Information). This allowed the cells to remain in suspension and to attach to the filaments of the scaffolds in a more homogenous manner. Around 20 to 30% of the total cell number in the cross section was found in each of the 1 mm subdivisions of the scaffold (**Fig. 2C and D**), regardless of the use of the MS-Dextran or the MS-Ficoll method, suggesting that both approaches successfully improved cell distribution along scaffold’s cross section. Furthermore, it is important to note that no cells were found at the bottom of scaffolds when seeded with the MS methods, contrary to what was observed in the case of CS scaffolds. It is plausible that the low cell number localized at the bottom filaments region, due to no cell sedimentation during MS methods, prevented the formation of a cell monolayer. Moreover, cells in suspension along the scaffold depth that did not attach to the filaments were washed away after immersing the scaffold in media (**Video S1**, Supplementary Information).

Direct comparison of our cell distribution results with previously published seeding techniques on AM 3D scaffolds is very challenging, as most of these studies have not shown the full characterization of cell distribution and sedimentation, but rather addressed the seeding efficiency and its direct effect on the tissue regeneration process. In one exception, the amount of DAPI nuclear staining was quantified in each of the slices subdividing the cross section of scaffolds with vertical pore size gradients [29]. Despite the positive effect of gradients on cell attachment in each of the scaffold subdivisions, still large amounts of cells accumulated at the bottom of the scaffolds, leading to a non-uniform cell distribution over the scaffold cross section,. Interestingly, the uniform cell distribution demonstrated by MS methods closely resembles a recently published report in which hMSCs were seeded in a polymeric scaffold by means of a cell-laden polyethylene glycol dithiothreitol hydrogel [37]. In this hybrid system, the cell carrier hydrogel was hydrolytically degraded after 7 days, time that cells used to populate the scaffold’s filaments attaining a good distribution. Importantly, when using our novel MS method, cells attached directly to the filaments without the means of hydrogel carriers. As observed in **Fig. 3**A, cells preferentially attached to the top curved surface of the scaffold filaments and, due to cell clustering and a larger number of cells attached, cell projections are not clear in the MS-methods when compared with the spindle-like shape that the cells seeded by the CS method show after 4h seeding. Nevertheless, only after 24h post-seeding cells in the MS scaffolds showed a clear elongated morphology, characteristic of migrating cells, and covered the whole surface of the filaments in contrast with the less crowded filaments on CS scaffolds (**Fig. 3**B).

**Fig. 3.**
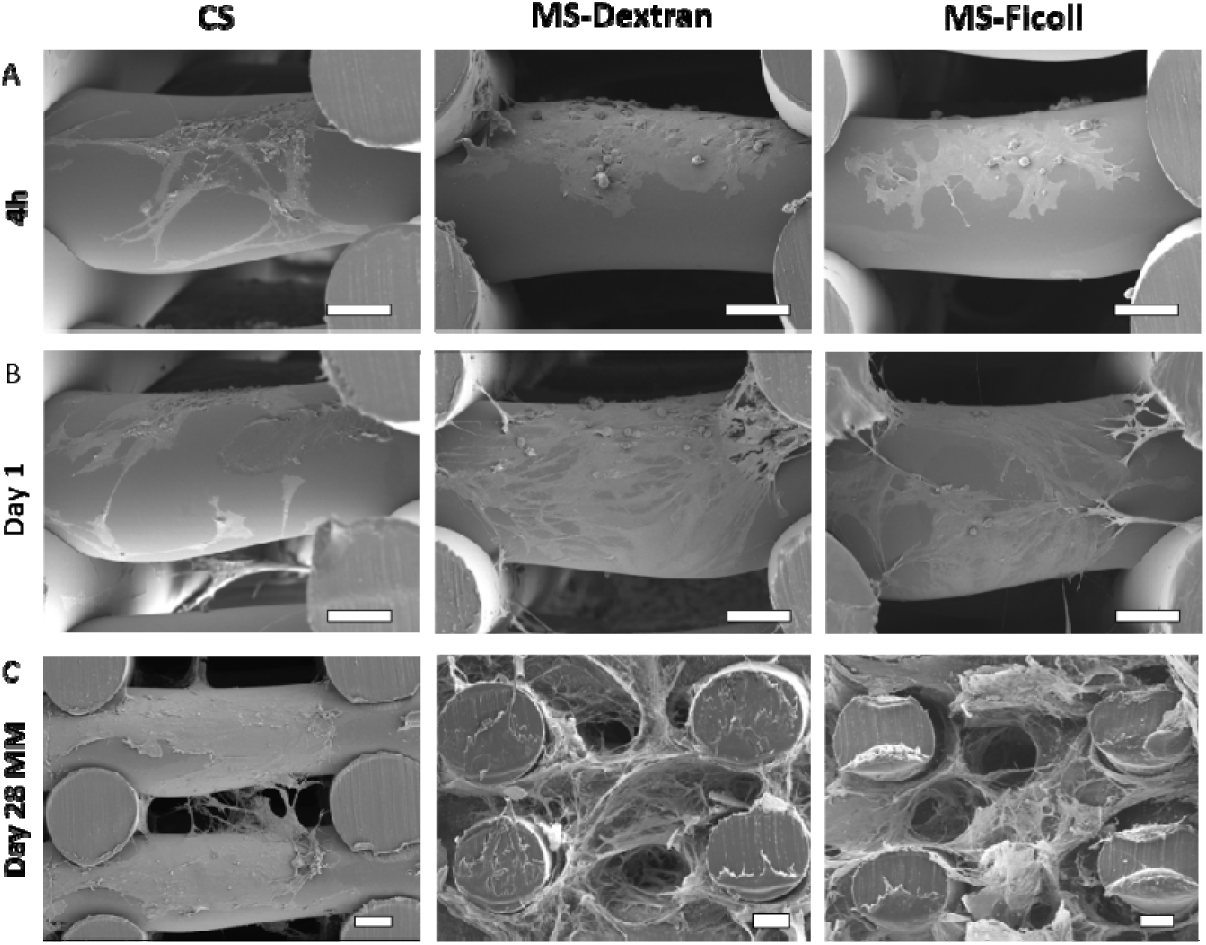
Higher cell coverage and matrix production on scaffolds seeded with the MS-methods. SEM micrographs of hMSCs on scaffolds seeded with the CS, MS-Dextran and MS-Ficoll methods after 4h, 1 day and 28 days of culture (21 days in MM) showing cell-scaffold interactions, cell coverage of scaffold filaments and matrix production. Scale bars 100 µm.

### 3.3. Macromolecules in scaffolds

In order to evaluate the potential presence of macroMs in the scaffolds after seeding, scaffolds were seeded with solutions containing fluorescently labeled dextran or Ficoll. The fact that macroMs were not found coating the scaffolds filaments and that cells sedimented on CS scaffolds pre-incubated with dextran or Ficoll based CM (**Fig. S3**, Supplementary Information), proved that the macroMs did not act as a cell adhesive coating in the scaffolds, but viscosity and density of the media were the only factors leading to cell attachment. Previous reports demonstrating reduced protein adsorption and cell attachment in dextran coated surfaces support these results [55]: it was claimed that dextran and Ficoll should be functionalized to render them cell adhesive [56].

Not surprisingly, after the 4h seeding, the only macroMs visible on the scaffold were found to co-localize with the cell cytoplasm (**Fig. 4**), suggesting that they were internalized by cells during the seeding period. Fluorescently labeled dextrans have been shown to enter the cells, also hMSCs [57, 58], by micropinocytosis and macropinocytosis and accumulate in lysosomal compartments [59, 60]. The fact that the fluorescence given by the macroMs was drastically reduced after 24h culture in CM, suggests that dextran and Ficoll were exocytosed towards the cell culture media. Moreover, since the media was renewed after 24h culture and every 2 days from then, the effect of the macroMs as macromolecular crowders influencing ECM deposition can be ruled out. Likewise, Ficoll can also be internalized by cells [52] and it is hypothesized to follow a similar endocytosis/exocytosis route as dextran, as both possess a similar chemical structure and size. It is important to realize that the macroMs observed in the scaffolds after 24h culture were found to colocalize only with dead cells (**Fig. S4**, Supplementary Information). As internalized macroMs remain in the lysosomes with no degradation or effect on cellular function before their exocytosis to the extracellular space, it is hypothesized that these cells were already dead before their incubation with macroMs. Thus, macroMs were able to penetrate their permeable membrane [61]. Cell death was visualized within the first minutes of hMSCs incubation with both macroMs based CM and CM (medium without macroMs) (**Fig. S5**, Supplementary Information). Hence, the presence of dead cells might be attributed to the low media-to-cell ratio during the 4h seeding and to the handling of cells prior to seeding (centrifugation and resuspension in a small volume of medium), which are unavoidable issues of the seeding process. Nevertheless, as demonstrated in **Fig. 5**, high cell viability was observed across the MS-Dextran and MS-Ficoll scaffolds over 7 days culture.

**Fig. 4.**
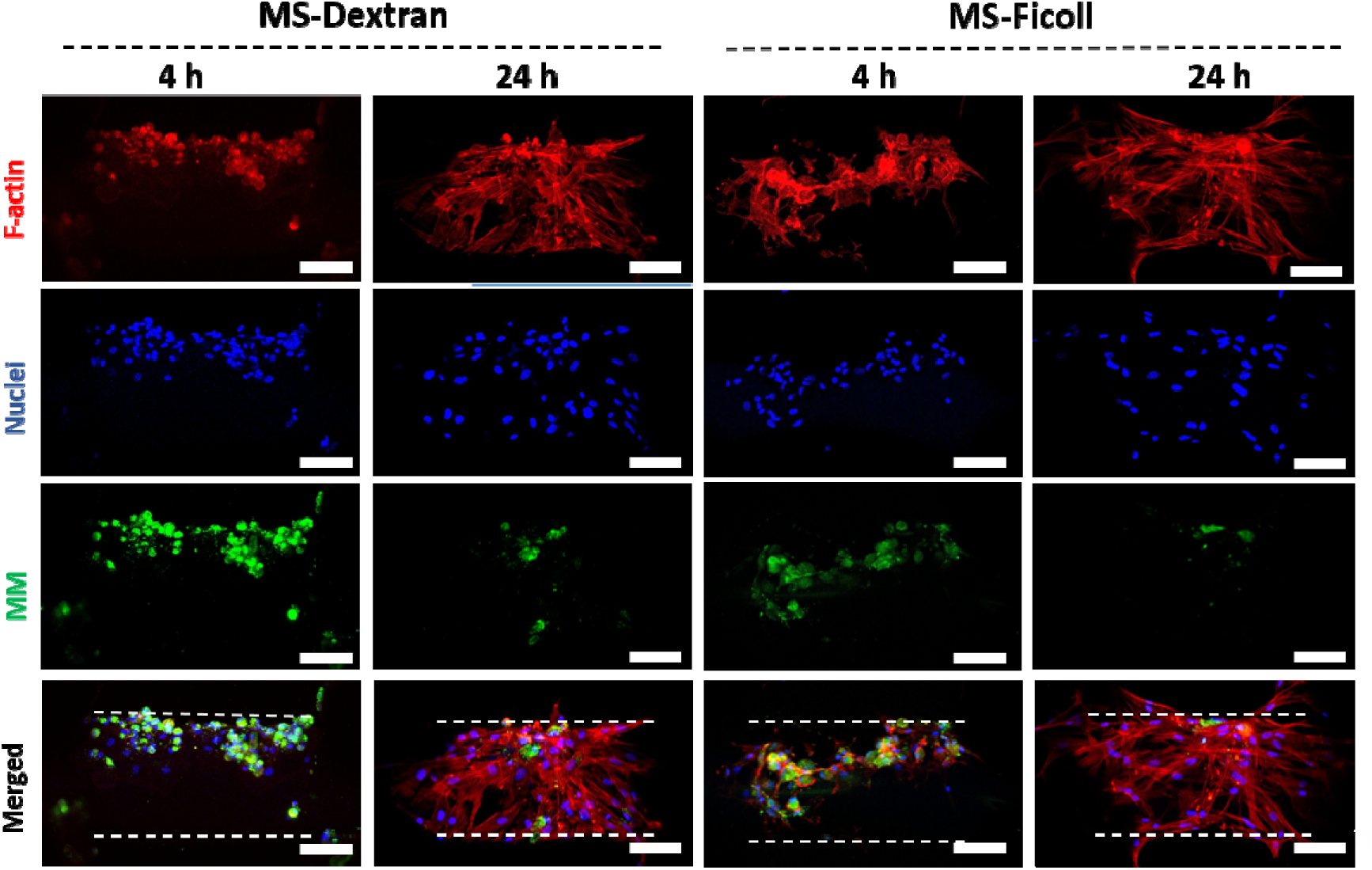
Initial retention (4h) and rapid elimination (24h) of macromolecules in cells attached to the scaffolds. Representative confocal microscopy images of hMSCS (F-actin) and macroM (FITC-labeled) on top of scaffold filament 4h and 24h post-seeding with the MS-Dextran and MS-Ficoll methods. Dash lines delimitating the scaffold filament. Scale bars 100 µm.

**Fig. 5.**
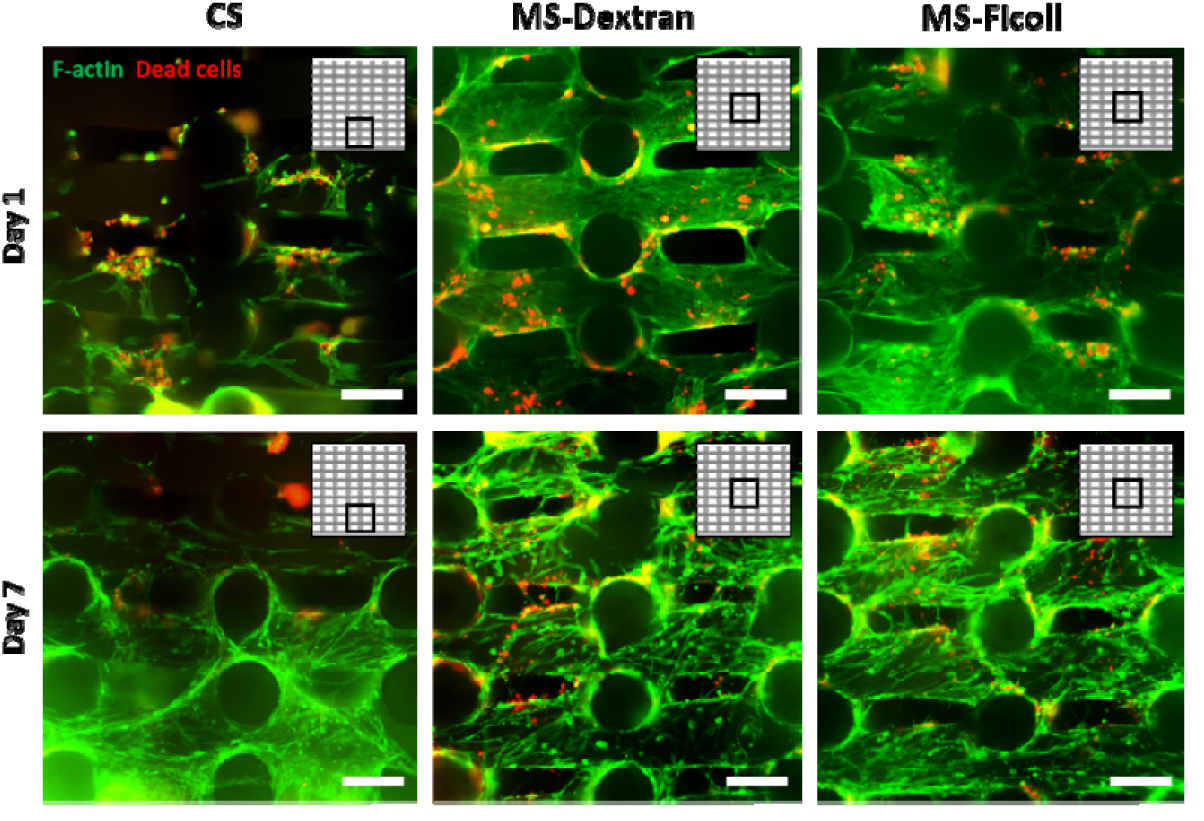
Cell viability is preserved after seeding with MS methods. Representative fluorescent images depicting cell viability (F-actin, green; dead cells, red) 1 day and 7 days post-seeding with the CS, MS-Dextran and MS-Ficoll methods. Inserts represent the imaged area in scaffold cross-section. Scale bars 250 µm.

### 3.4. Osteogenic differentiation on MS scaffolds

To fully assess the functionality of the novel MS methods to seed AM 3D scaffolds for bone regeneration, hMSCs potency and ability to undergo osteogenic differentiation after seeding was evaluated. Preliminary 2D experiments confirmed that proliferation and mineralization were not affected when cells pre-incubated in macroMs solutions and seeded in 2D (**Fig. S6,** Supplementary Information). These initial positive results allowed us to further study hMSCs response to macroMs seeding in 3D. Although the MS methods had a clear effect in cell distribution (**Fig. 2**), no statistical differences were found in the total number of cells attached to the scaffolds after seeding when compared with the CS method (**Fig. 6**A). It is plausible that the sedimented cells which attached to the bottom of the CS scaffolds, as previously mentioned, contributed to the relatively higher seeding efficiency values than expected (∼55%) for the high amount of cell settling observed, making it comparable to the MS methods seeding efficiency (∼80% and ∼75%, for MS-Dextran and MS-Ficoll, respectively). In 2D, high cell seeding density (cells*cm^-1^) has been demonstrated to direct and control early and late stage osteogenic markers and the temporal expression of MSCs genes towards osteogenesis [62]. However, cell density coupled to an optimum distribution of cells occupying the whole scaffold surface area are the parameters that will ultimately favor osteogenesis in a 3D environment. Therefore, cell number alone is not a primary factor and “efficiency” is defined by both cell density and cell distribution when culturing 3D scaffolds.

**Fig. 6.**
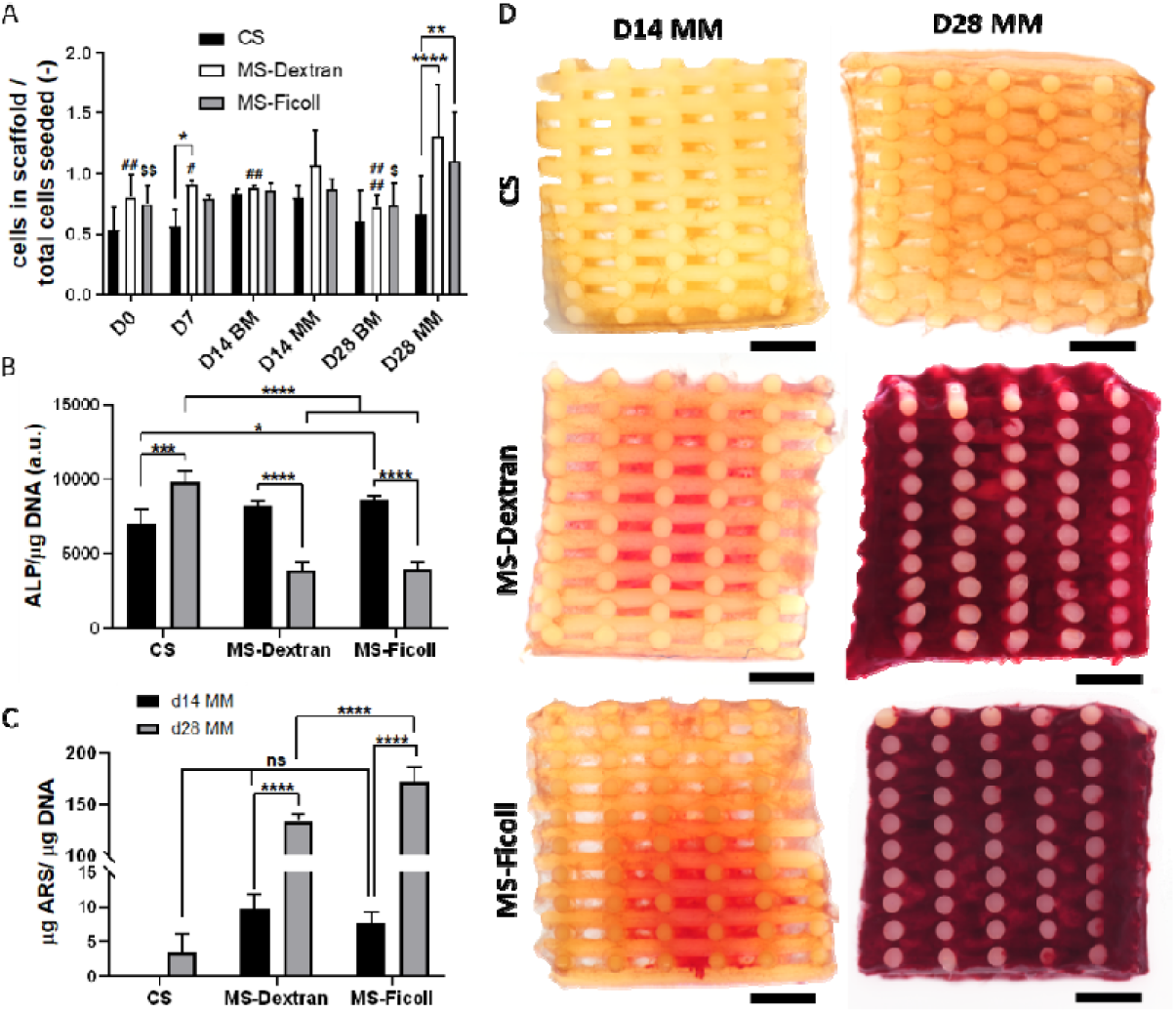
MS method does not hinder osteogenic differentiation of hMSCs: Evaluation of the osteogenic differentiation potential of hMSCS seeded with the MS-based methods compared to CS. (a) Cell seeding efficiency and cell number progression over 28 days of culture in BM and MM on scaffolds seeded with the CS, MS-Dextran and MS-Ficoll methods. Statistical significance performed using two-way ANOVA with Tukey’s multiple comparison test (*#$ p < 0.05; **##$$ p < 0.01; ****####$$$$ p < 0.0001; * for comparisons among seeding methods each time point; # for comparisons with D28 MM among MS-dextran; $ for comparisons with D28 MM among MS-Ficoll). (b) Comparison of ALP activity at day 14 and day 28 of culture (7 and 21 days in MM, respectively) of hMSCs when seeded with the three different methods. (c) Quantification of the alizarin red S staining extracted from scaffolds after 14 and 28 days of culture (7 and 21 days in MM, respectively), normalized to cell number. Statistical significance performed using two-way ANOVA with Sidak’s multiple comparison test (*p < 0.05; ***p < 0.001; ****p < 0.0001, for comparisons among seeding methods each time point or among time points each seeding method) (d) Stereomicroscopy images of scaffold cross sections stained with alizarin red S after 14 and 28 days of culture (7 and 21 days in MM, respectively). Scale bars 1 mm.

Looking at cell proliferation (**Fig. 6**A), constant cell numbers were observed in scaffolds seeded with all the three methods over time. Due to higher chance of cell attachment when seeded with the MS methods, cells were confluent occupying the scaffold filaments from the beginning of the culture, as observed in the scaffold cross section (**Fig. 2** and **Fig. 3**B). This saturation phenomenon may explain the lack of proliferation on these scaffolds over 28 days in BM [63]. Cell number increased only at day 28 in MM on MS scaffolds, compared with earlier time points and with CS scaffolds. This is possibly due to the extensive ECM production on MS scaffolds that enabled cell proliferation also within the volume of the deposited matrix in scaffolds’ pores at this time point. This was further confirmed with the SEM images in **Fig. 3**C demonstrating the presence of cells both on the scaffold filaments and within pores’ volume along with the formation of a dense ECM. On the contrary, ECM density was found to be much lower in CS scaffolds, which correlates to the low cell proliferation values over the whole culture period (**Fig. 6**A).

Alkaline phosphatase activity (ALP) was measured after 14 and 28 days of culture, corresponding to 7 and 21 days culture in MM (**Fig. 6**B). ALP is an early stage enzyme expressed in bone development and its upregulation precedes the upregulation of late osteogenic genes such as osteocalcin [64, 65]. No significant difference was observed in ALP activity at day 14 regardless of the seeding method. However, ALP activity was significantly lower at day 28 between cells in scaffolds seeded with CS and those seeded with MS methods. Interestingly, ALP activity increased from day 14 to day 28 in cells seeded with CS method. Concordantly with our results, a peak of ALP activity after 28 days of culture on scaffolds seeded with hMSCs via the CS method has previously been reported [66, 67]. In contrast, ALP decreased from day 14 to day 28 in cells seeded with MS methods despite the macroM type. This is likely due to the aforementioned differences in cell number coupled to their distribution profile within the scaffolds. Decreasing ALP activity over time, which enabled the development of the bone matrix, has previously been observed in scaffolds with high cell seeding numbers, and therefore possibly better cell distribution [68].

To gain more insight into the potential of osteogenic differentiation of hMSCs in scaffolds seeded with our novel method, matrix mineralization was evaluated using alizarin red S staining to visualize calcium deposits after 7 and 21 days in MM (day 14 and day 28, respectively). Scaffolds cultured in BM were used as controls to ensure the inert behavior of the macroMs in the mineralization process (**Fig. S7**A, Supplementary Information). Intriguingly, as it is shown in **Fig. 6**D, calcium deposition on the cross section of MS-scaffolds was observed from day 14 (7 days in MM), and by day 28 these were distributed over the whole scaffold visualized by the dense red color homogeneously covering the cross section and outer surface of the scaffolds (**Fig. S7**B, Supplementary Information). Quantification of the staining (**Fig. 6**C) confirmed that calcium depositions increased ∼10-15 fold within these two time points. Moreover, statistical differences among the amount of calcium deposited by cells seeded by the MS-dextran and the MS-Ficoll methods were evident when quantified. However, no clear hypothesis has been drawn to explain this result, since cell number at the time point are comparable in both scaffolds and cell viability was also preserved in both cases. As expected from the ALP activity results, no ARS staining was observed in CS scaffolds at day 14, and only a small amount of calcium deposits was quantified after 28 days culture, which was comparable to the mineralization in MS scaffolds at day 14.

Ultimately, the expression of relevant osteogenic genes on cells seeded with the MS-methods was evaluated and compared to the CS-method. Notably, no significant differences were found in the expression of runt-related transcription factor (RUNX2), collagen I, osteocalcin and osteonectin among the different groups at day 14 (7 days in MM), suggesting that the seeding with macroM did not alter the early stage of hMSCs differentiation (**Fig. 7**). Importantly, the gene encoding RUNX2, which is an essential transcription factor regulator of hMSCs differentiation into the osteogenic lineage [69] was upregulated on cells seeded with the MS-Dextran and MS-Ficoll at day 28 (21 days in MM) when compared to CS method (**Fig. 7**A). Similarly, collagen I expression, one of the major bone ECM components, was higher in MS-Ficoll samples at this time point (**Fig. 7**B). Osteocalcin, one of the major bone non-collagenous proteins with the ability to bind bone hydroxyapatite [70], was more expressed in MS-Dextran with respect to CS (**Fig. 7**C). On the other hand, no significant differences were found in the osteonectin encoding gene expression among time points or seeding methods (**Fig. 7**D). However, this calcium binding non-collagenous protein has been found to be expressed in other non-mineralizing tissues, therefore lacks specificity [71]. The upregulation of some of the aforementioned protein encoding genes in cells seeded with MS-methods, complements the abovementioned ALP activity and calcium deposition results towards the confirmation that there is either a delay in the progression of osteogenesis in cells seeded with the CS method, or an acceleration in cells seeded with the MS methods, due differences in homogeneity in cell distribution. While longer culture periods of CS scaffolds in MM might lead to comparable results among different seeding methods [72], the possibility of speeding the osteogenic process of hMSCs by MS-methods is desired for both the in vitro and in vivo applications of the scaffolds.

**Fig. 7.**
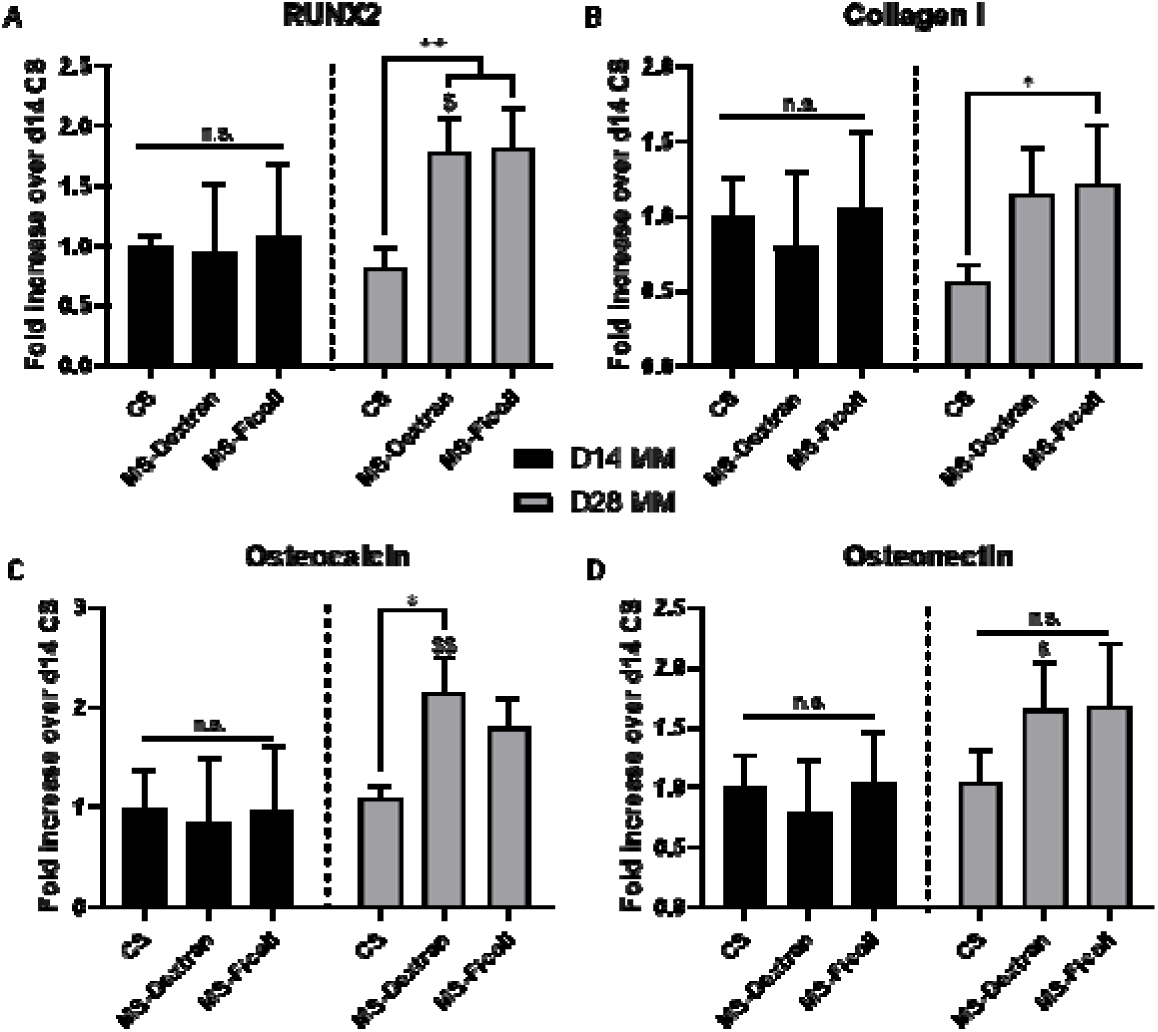
Gene expression of hMSCs after 14 and 28 days of culture (7 and 21 days in MM, respectively) comparing the three different seeding methods. (a) RUNX2, (b) collagen 1, (c) osteocalcin and (d) osteopontin fold change expression values are relative to CS d14 MM. Data presented as average ± s.d. and statistical significance performed using two-way ANOVA with Sidak’s multiple comparison test (*$ p < 0.05; **$$ p < 0.01; * for comparisons among seeding methods each time point; $ for comparisons among time points each seeding method).

## 4. Conclusions

Cell seeding on 3D AM scaffolds made out of bioinert synthetic polymers with interconnected macroporosity still remains a challenge. The aim of this study was to develop a simple and reliable seeding technique to improve cell distribution upon static seeding valid for all 3D AM scaffolds regardless of their material properties and architecture. This was achieved by controlling the settling velocity of cells by altering the viscosity and density of the seeding media via the addition of two different macromolecules, Dextran and Ficoll. Compared to scaffolds seeded with conventional static techniques, a homogeneous cell distribution was attained with direct cell attachment to the scaffold filaments. Importantly, macromolecules did not affect viability or osteogenic differentiation of hMSCs and their complete removal was demonstrated after the seeding period. Intriguingly, the improved cell distribution from the macromolecules based seeding led to enhanced mineralization and bone matrix maturation compared to conventional seeding. This newly proposed seeding method is simple and reproducible and has the potential to improve the long-term functionality of *in vitro* and *in vivo* constructs for tissue engineering applications.

## Data availability statement

The raw/processed data required to reproduce these findings cannot be shared at this time as the data also forms part of an ongoing study.

## Acknowledgements

We are grateful to H2020-NMP-PILOTS-2015 (GA n. 685825) for financial support.

## Appendix A. Supplementary data

Supplementary data related to this article can be found at…

