## Supplementary Information for "Improving cell distribution on 3D additive manufactured scaffolds through engineered seeding media density and viscosity"

**Table S1.** Primer sequences used for q-PCR. B2M: beta-2-microglobulin; RUNX2: runt-related transcription factor; COL1A1: collagen 1 alpha 1; OCN: osteocalcin: SPARC: osteopontin.

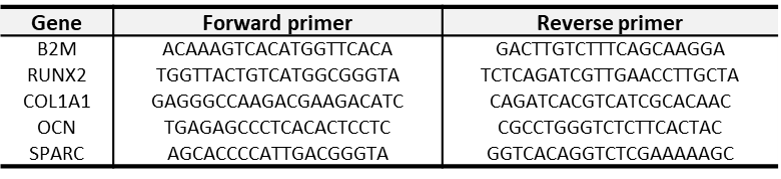

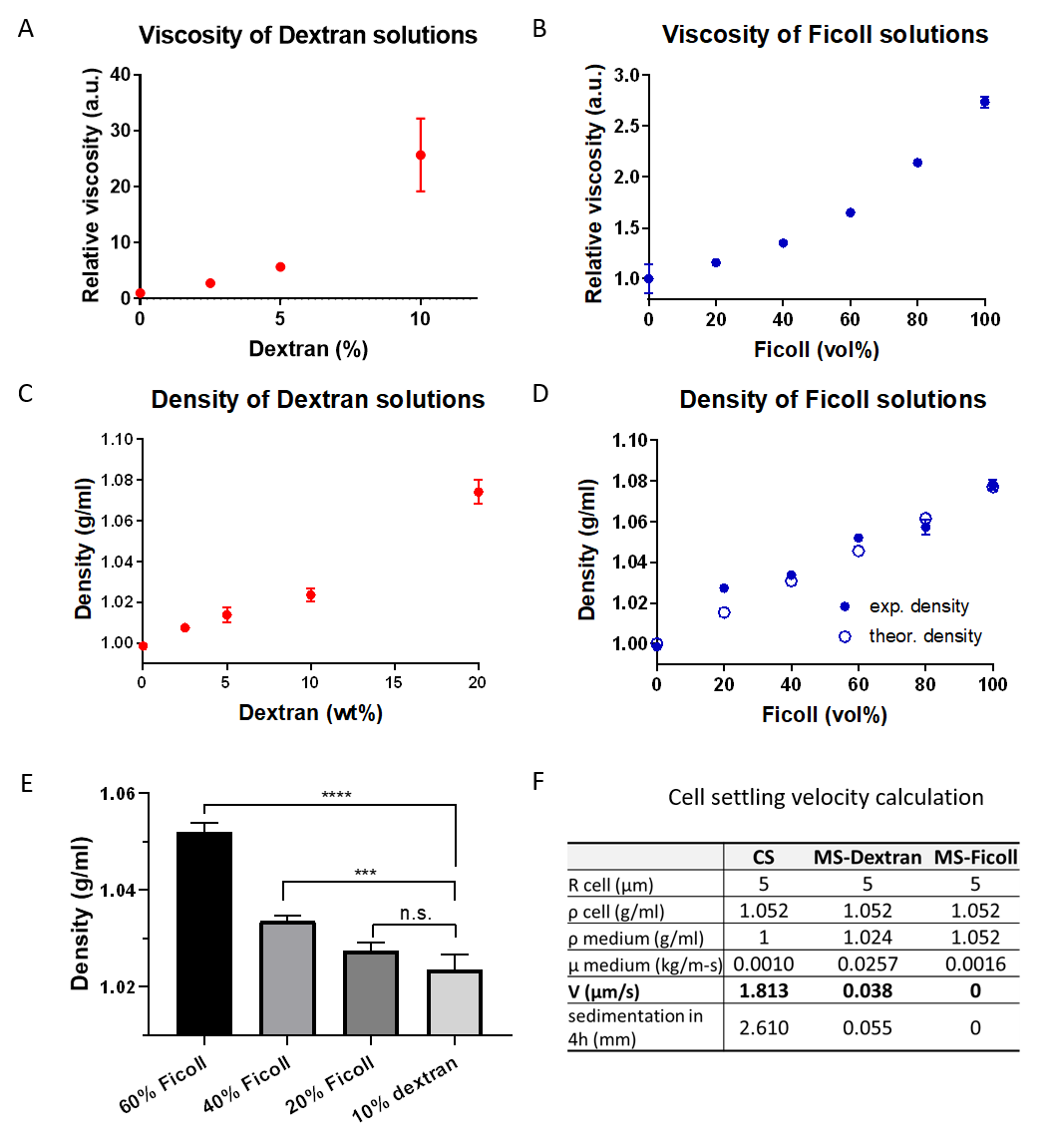

**Fig. S1.** (a, b) Relative viscosity and (c, d) density of cell culture media containing different concentrations of dextran and Ficoll. (e) Comparison of density values of media containing specific dextran and Ficoll concentrations. (f) Cell settling velocity in CS, MS-Dextran and MS-Ficoll methods.

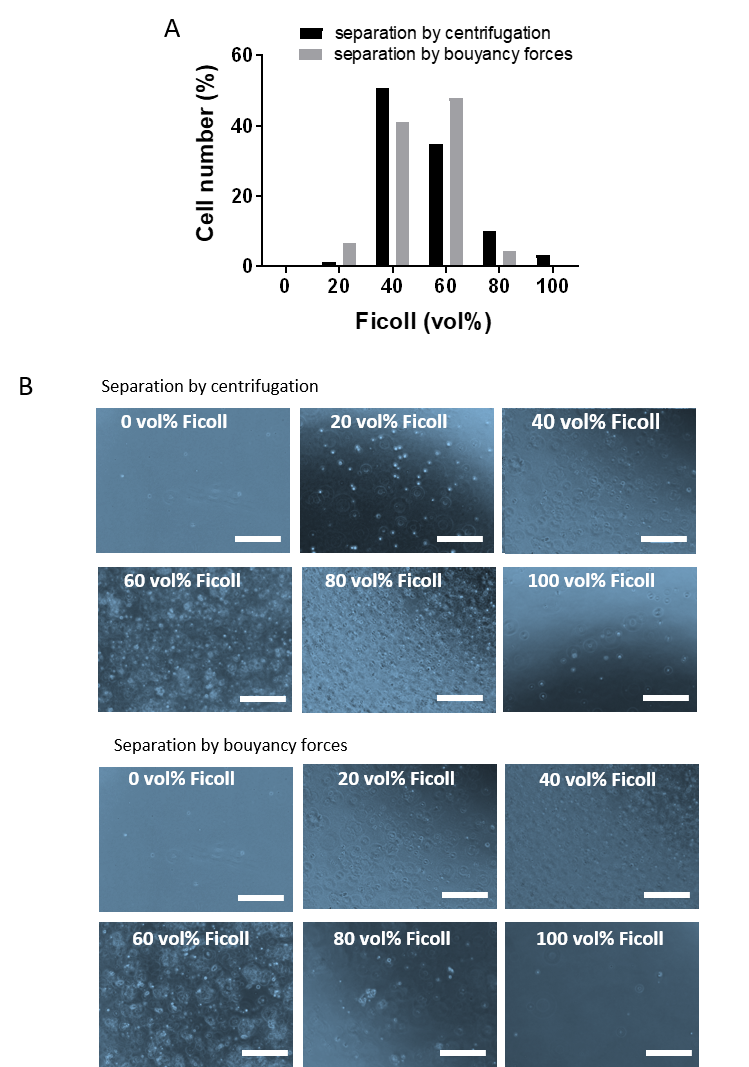

**Fig. S2.** (a) Quantification of cell number in each layer of the Ficoll gradient used to determine hMSCs population density by separation by centrifugation or by separation by buoyancy forces. (b) Representative images of the cells found in each layer after each separation method. Scale bars 200 µm.

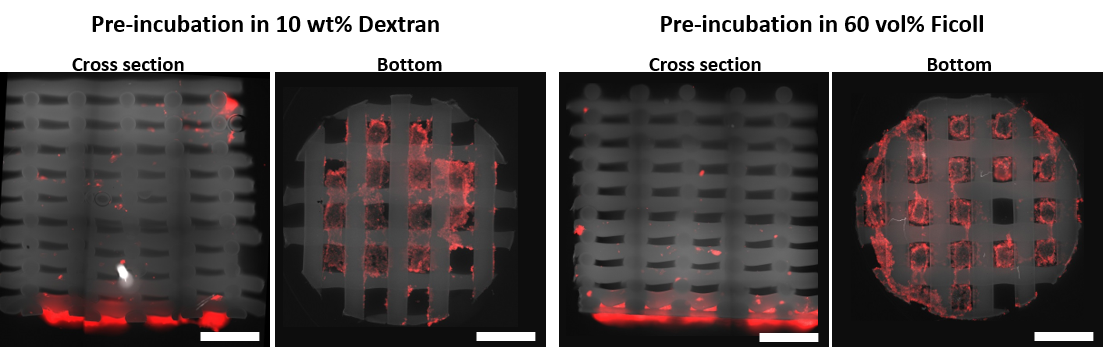

**Fig. S3.** Fluorescence microscopy images of hMSCs in the cross sections and bottom lids of scaffolds pre-incubated with dextran and Ficoll based solutions and seeded with the CS method. Scale bar 1 mm.

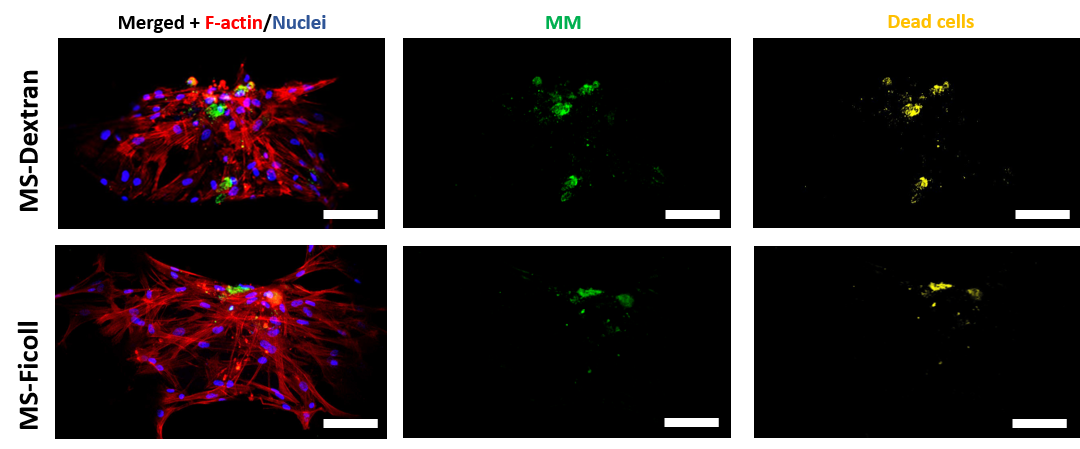

**Fig. S4.** Representative confocal microscopy images of hMSCS (F-actin and nuclei), macroMs (FITC-labeled) and dead cells on top of scaffold fiber 24h post-seeding with the MS-Dextran and MS-Ficoll methods. Scale bars 100 µm.

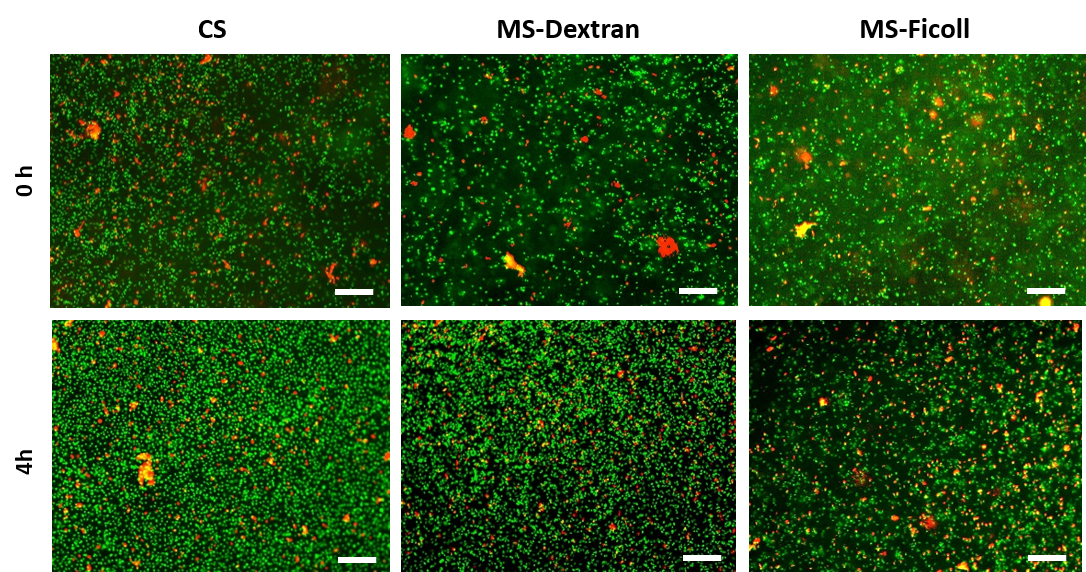

**Fig. S5.** Representative fluorescent images depicting hMSCs viability (live, green; dead, red) right after their resuspension (0 h) and after 4h incubation in macroM based solutions, as compared to control (CS). Scale bars 250 µm.

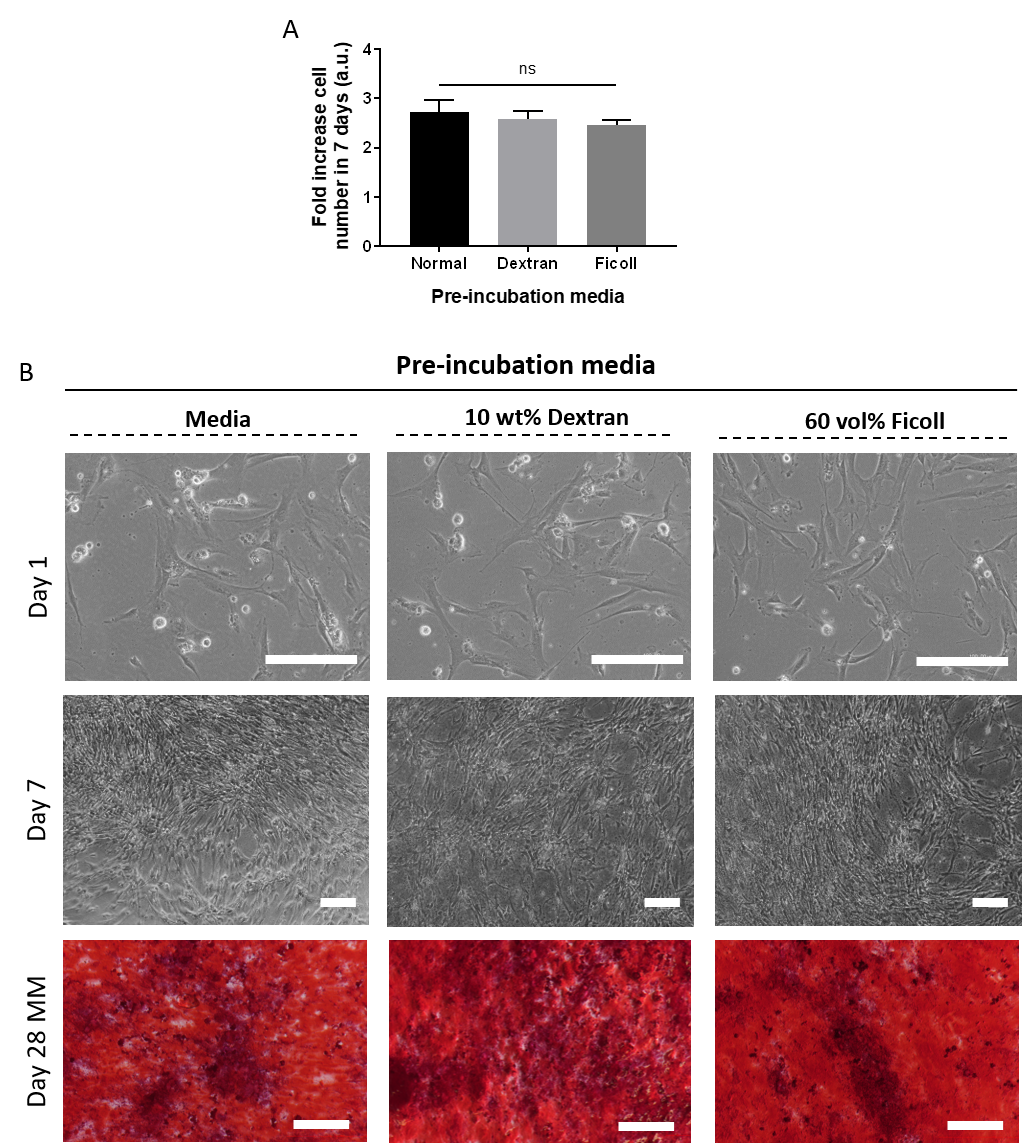

**Fig. S6.** Cell culture and differentiation in 2D after hMSCs incubation in macroM based solutions during 4h mimicking the scaffold seeding conditions. (a) Fold increase in cell number after 7 days of culture in BM and pre-incubation in normal, dextran or Ficoll based CM. (b) Representative brightfield images of cells pre-incubated in normal, dextran or Ficoll based CM after 1 and 7 days of culture in BM. (c) Stereomicroscopy images of cell monolayers stained with alizarin red S after 28 days of culture (21 days in MM) and pre-incubated in normal, dextran or Ficoll based CM. Scale bars 250 µm.

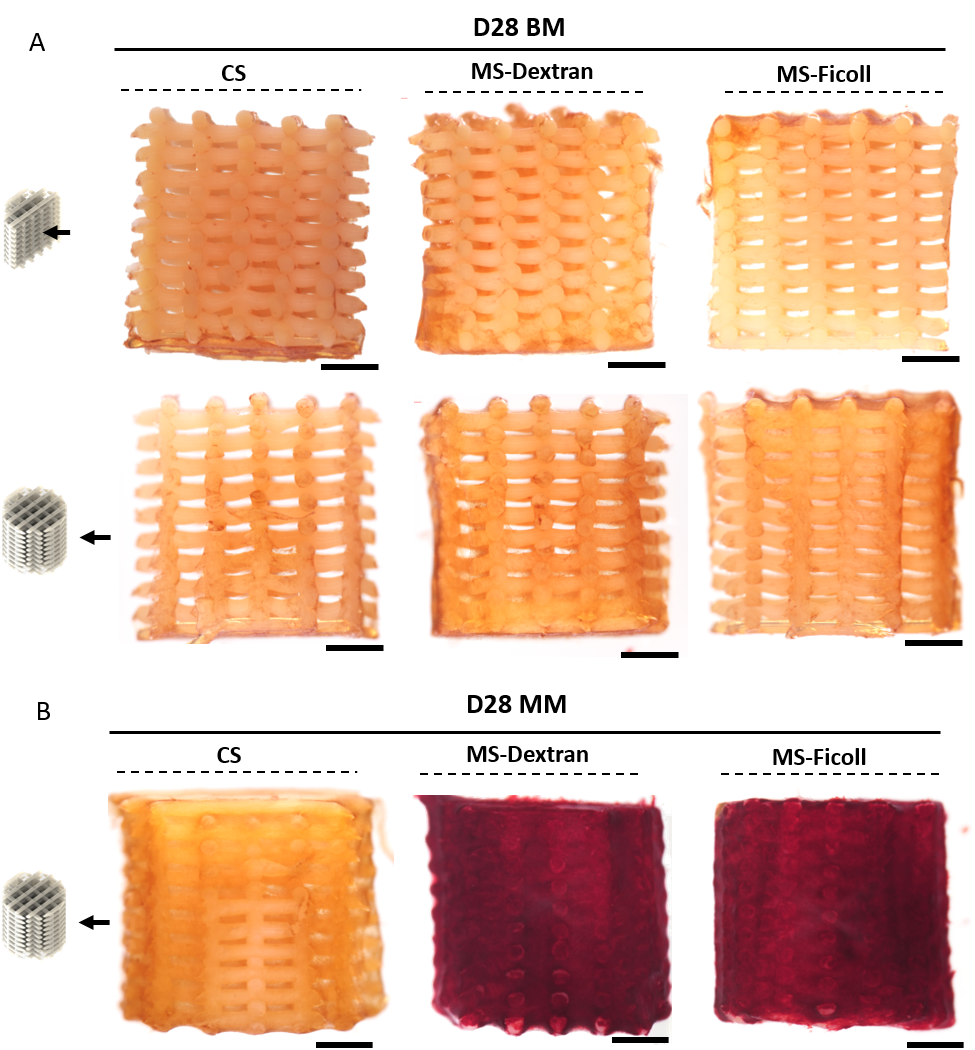

**Fig. S7.** Stereomicroscopy images of alizarin red S stained scaffolds. (a) Cross- and outer sections after 28 days of culture in BM and (b) outer sections after 28 days of culture (21 days in MM). Scale bars 1 mm.

**Video S1**. MS-Dextran scaffold: after seeding, scaffold is placed in media and cells that did not attach to the filaments are washed away.
